# Lipid droplet targeting of ABHD5 and PNPLA3 I148M is required to promote liver steatosis

**DOI:** 10.1101/2024.10.03.616525

**Authors:** Grace Teskey, Nivedita Tiwari, Andrew J. Butcko, Amit Kumar, Anu Yadav, Yu-ming M. Huang, Christopher V. Kelly, James G. Granneman, James W. Perfield, Emilio P. Mottillo

## Abstract

The storage and release of triacylglycerol (TAG) in lipid droplets (LDs) is regulated by dynamic protein interactions. α/β hydrolase domain-containing protein 5 (ABHD5; also known as CGI-58) is a membrane/LD bound protein that functions as a co-activator of Patatin Like Phospholipase Domain Containing 2 (PNPLA2; also known as Adipose triglyceride lipase, ATGL) the rate-limiting enzyme for TAG hydrolysis. The dysregulation of TAG hydrolysis is involved in various metabolic diseases such as metabolic dysfunction-associated steatotic liver disease (MASLD). We previously demonstrated that ABHD5 interacted with PNPLA3, a closely related family member to PNPLA2. Importantly, a common missense variant in PNPLA3 (I148M) is the greatest genetic risk factor for MASLD. PNPLA3 148M functions to sequester ABHD5 and prevent co-activation of PNPLA2, which has implications for initiating MASLD; however, the exact mechanisms involved are not understood. Here we demonstrate that LD targeting of both ABHD5 and PNPLA3 I148M is required for the interaction. Molecular modeling demonstrates important resides in the C-terminus of PNPLA3 for LD binding and fluorescence cross-correlation spectroscopy demonstrates that PNPLA3 I148M greater associates with ABHD5 than WT PNPLA3. Moreover, the C-terminus of PNPLA3 is sufficient for functional targeting of PNPLAs to LD and the interaction with ABHD5. In addition, ABHD5 is a general binding partner of LD-bound PNPLAs. Finally, PNPLA3 I148M targeting to LD is required to promote steatosis in vitro and in the liver. Overall results suggest that PNPLA3 I148M is a gain of function mutation and that the interaction with ABHD5 on LD is required to promote liver steatosis.

## INTRODUCTION

The storage and hydrolysis of triacylglycerol (TAG) is regulated by dynamic protein-protein interactions on the surface of lipid droplets (LDs) in key metabolic tissues such as adipose tissue and liver. α/β hydrolase domain-containing 5 (ABHD5), an LD protein and lipase co-activator, is a critical determinant of liver TAG levels. As such, genetic deletion of ABHD5 in mice and point mutations in humans promote MASLD (1, 2). ABHD5 co-activates Patatin-Like Phospholipase Domain Containing 2 (PNPLA2; widely known as Adipose Triglyceride Lipase, ATGL), the rate limiting TAG hydrolase to greatly increase its activity and TAG mobilization in the form of free fatty acids (FFA) (3). Importantly, increasing genetic and biochemical evidence suggest that the dysregulation of these protein interactions can impair the balance between TAG storage and mobilization and promote lipid accumulation in the liver that leads to metabolic dysfunction–associated steatotic liver disease (MASLD) (4–6).

We recently discovered the interaction between ABHD5 and PNPLA3, a closely related family member to PNPLA2 (6). The ABHD5-PNPLA3 interaction is important for metabolic health, as a common genetic variant of PNPLA3 (rs738409; minor allele frequency ∼ 35% (7)), I148M, is the most prevalent genetic risk factors for the development of MASLD (8, 9). ABHD5 interacts with WT PNPLA3 on the endoplasmic reticulum (ER) and LDs, while the interaction with PNPLA3 I148M occurs mostly on LDs (6). Importantly, PNPLA3 I148M sequesters ABHD5 away from PNPLA2 leading to steatosis (6) and modifies TAG metabolism in an ABHD5-dependent manner (4, 6).

While the interaction between ABHD5 and PNPLA3 describes a potential mechanism by which the I148M variant causes MASLD, a better understanding of the biochemical basis of the interaction is required to develop novel therapeutics. Here we demonstrate that that localization of ABHD5 and PNPLA3 I148M to LDs is required for their interaction. The C-terminal domain of PNPLA3 is sufficient to confer LD targeting and promote the interaction with ABHD5 and molecular dynamic simulations demonstrate how residues in the C-terminus of PNPLA3 confer LD binding. Moreover, preventing PNPLA3 I148M from targeting to LDs blocks the interaction with ABHD5 and the ability to promote hepatosteatosis. Overall, these data improve our understanding of the mechanism by which PNPLA3 I148M causes steatosis and could lead to novel therapies to treat MASLD.

## EXPERIMENTAL PROCEDURES

### Plasmids and Cloning

Human PNPLA3 I148M (hPNPLA3 I148M) lipid droplet mutants were generated using PCR Overlap Extension PCR using hPNPLA3 I148M as a template with hPNPLA3 I148M ^370^QAAA^373^ forward and reverse primers, and hPNPLA3 I148M ^370^AAEE^373^ forward and reverse primers (Refer to Supplementary Table 1).

After amplification the PCR products were cloned in frame with the N-terminus of *gaussia* luciferase (GlucN) using HindIII and AgeI sites as previously described.(6) EYFP tagged version were made by replacing the GlucN fragment with EYFP using AgeI and NotI restriction sites. hPNPLA6, hPNPLA7, hPNPLA8 and hPNPLA9 were amplified from U20S or 293T cDNA using following primers and cloned with in frame with GlucN vector using the following primers and restriction sites: Pnpla6 Forward and reverse using NheI and BamH1; Pnpla7 forward and reverse using NheI and HindIII; Pnpla9 forward and reverse using HindIII and AgeI (Refer to Supplementary Table 1 for primer sequences). Human ABHD5-ECFP was created by subcloning the ECFP fragment onto the C-terminus of ABHD5 using AgeI and NotI restriction sites. Mouse PNPLA5 was PCR amplified from a cDNA clone from Open Biosystems (Cat No. MMM1013-211693311).

hPNPLA3 WT-EYFP and hPNPLA3 I148-EYFP plasmids were created by subcloning EYFP onto the C-terminus of PNPLA3 using AgeI and NotI restriction sites. Fusions of PNPLA3 and PNPLA4 were created by PCR amplification of the C-terminus of hPNPLA3 (320–481) fusion using hPNPLA3 I148M-AAEE-EYFP and hPNPLA3-I148M-EYFP plasmids as a template for amplification with the following primers: forward hPNPLA3 320 and reverse hPNPLA3 481. PCR products were cloned into hPNPLA4-Gluc N and hPNPLA4-EYFP vector using the AgeI restriction site. All PCR generated plasmids were confirmed with Sanger sequencing.

AAV vectors for the expression of EGFP pENN.AAV.TBG.PI.EGFP, human PNPLA3 (hPNPLA3) pENN.AAV.TBG.PI.hPNPLA3 and human PNPLA3 I148M pENN.AAV.TBG.PI.hPNPLA3 I148M were produced as AAV8 serotype and obtained from Vector Biolabs (Malvern, PA). PNPLA3 I148_AAAE_ was PCR amplified from the hsPNPLA3 I148M AAEE-GlucN template and cloned into the pENN.AAV.TBG.PI.ffLuciferase into the KpnI and MluI restriction sites and AAVpENN.AAV.TBG.PI.ffLuciferase was a gift from James M. Wilson (Addgene plasmid # 105538; http://n2t.net/addgene:105538; RRID:Addgene_105538). The AAV for hsPNPLA3 I148M AAEE was produced at the University of Michigan Vector core as an AAV8 serotype.

### Cell culture and Transfections

COS7, U2OS (ATCC #HTB-96) and HEK 293A (ATCC #CRL-1573) cells were cultured in DMEM High Glucose (Fisher Scientific, SH30243.01) supplemented with 10% fetal bovine serum (Fisher Scientific, SH3039603) and 5% penicillin/streptomycin (Fisher Scientific, SV30010). Cells were incubated at 37°C and 5% CO_2_.

HEK 293A cells were transfected in preparation for the luciferase assay. Cells were seeded in a 24 well plate at a density of 40,000 cells/well and allowed to adhere overnight. Cells were transfected using Lipofectamine 3000 Transfection Reagent (Thermo Fisher Scientific,

#L300015), with 0.5µg of total DNA/well, 1µL of P3000 reagent/well and 1.5µL of Lipofectamine 3000 reagent. The DNA-lipid complex was prepared in Opti-MEM reduced serum medium (Thermo Fisher Scientific, #11058021) The transfected cells were incubated for 6 hours and the media was removed from the cells and replaced with complete DMEM growth medium with or without 0.2 mM oleic acid complexed with bovine serum albumin (BSA).

U2OS or COS7 cells were prepared for fluorescent microscopy by seeding on coverslips in a 6 well plate at a density of 300,000 cells/well and allowed to adhere overnight. Cells were transfected using GenJet Plus DNA In Vitro Transfection Reagent (SignaGen Laboratories, #SL100499), with 1µL of total DNA/well and 3µL of GenJet Plus transfection reagent/well. DNA complexes were made in serum-free. Following transfection cells were treated as above for 293A cells.

### Luciferase Assay

*Gaussia* luciferase (Gluc) activity for protein complementation assays (PCA) was measured as previously described (10). In general, transfected cells were washed with PBS then lysed with 150µL/well of Intracellular Buffer (10mM HEPES, pH 7.3, 140mM KCl, 6mM NaCl, 1mM MgCl2, 2mM EGTA) and three freeze/thaw cycles. 100µL of cell lysate from each well was loaded into a 96 well plate (Greiner Bio-One, #655098). Luminescence was quantified using a Clariostar luminometer after the addition of 100µL of 40µM coelenterazine substrate (Goldbio, #CZ10) in PBS.

### Fluorescence Microscopy and Fluorescence Energy Transfer (FRET)

Transfected cells were washed with PBS and fixed with 4% paraformaldehyde. When required for the experiment, cells were stained with HCS LipidTOX Red (Thermo Fisher Scientific, #H34476) diluted 1:1000 in PBS for 30 minutes at room temperature. Images were acquired using an Olympus IX-81 microscope equipped with a spinning disc confocal unit and filter sets as previously described (11). Microscope control and data acquisition was performed using CellSens Dimensions software. Fluorescent signal from YFP, ECFP and mCherry was acquired at a magnification of 60X (1.2 NA) plan apo water immersion lens and a Hamamatsu ORCA cooled CCD camera. FRET was performed using the three-filter method and the net FRET (nFRET) was calculated using the FRET extension of the CellSens Dimensions software.

### Animal Studies

PNPLA3 knockout mice were provided by Eli Lilly and Company. All mice were housed at 24 °C ± 2 °C with a 12 h light:12 h dark cycle in an American Association for Laboratory Animal Care approved animal facility at Wayne State University. All protocols involving animals were approved by the Institutional Animal Care and Use Committee of Wayne State University and followed the National Institutes of Health Guide for the Care and Use of Laboratory Animals. Male mice between 8 and 12 weeks of age were tail vein injected with 1X10^6^ genome copies (gc) of AAV8 serotype. Three days later mice were fed a high sucrose diet (74% kcal from sucrose, MP Biomedicals) for 4 weeks. Mice were anesthetized with isoflurane and liver tissue was processed for fractionation as previously described (6) with minor modification with centrifugation in a Beckman Coulter 22R Refrigerated Microcentrifuge containing Beckman Microfuge Swinging Bucket Rotor (S241.5) for 30 min at 18000 g. The pellet containing membranes and mitochondria was solubilized with 1% SDS.

### Triacylglycerol Quantification

Neutral lipids were extracted from liver via the method of Folch (12) and the amount TAGs were quantified with a Serum Triglyceride Determination Kit (Sigma) on a BioTek Synergy plate reader at 540 nm absorbance and normalized to wet tissue weight.

### Immunoblot analysis

Immunoblot analysis was performed as previously described (6). Briefly, total-cell proteins were extracted with RIPA buffer (Teknova) with a protease inhibitor tablet (Pierce). Samples were run on Mini-Protean TGX Gels (BioRad), transferred to PVDF membranes and blocked in 5% non-fat milk for 1 h. Membranes were probed for 1 hr or overnight as indicated by the antibody manufacturer. The following antibodies were used: rabbit anti-Gaussia luciferase (Nanolight, cat. no. 401P) rabbit anti-β-actin HRP conjugate (Cell Signaling cat. no. 5125S), Mek1/2 (Cell Signaling cat. No. S8727), AIF (Cell Signaling cat. No. S5318), chicken anti-PLIN2 (Abcam cat. No. ab37516) and human PNPLA3 (R&D Systems, cat. No. AF5208).

### Coarse grained molecular dynamics (CGMD) and Gaussian accelerated molecular dynamics (GaMD) simulations

Since the experimental PNPLA3 structure is unknown, all-atom (AA) PNPLA3 model containing 481 residues was built using AlphaFold2 (13). The LD was modeled as a bilayer composed of palmitoyl-2-oleoyl-glycero-3-phosphocholine (POPC), Dioleoylphosphatidylethanolamine (DOPE), and 1-stearoyl-2-arachidonoyl-phosphatidylinositol (SAPI) lipids in a ratio of 88:37:10, with a 4 nm thick TAG layer sandwiched between the bilayer. The bilayer was assembled using CHARMM-GUI (14) and the TAG layer was constructed using PACKMOL (15). Each system was solvated with TIP3P water molecules (16), with a 15 Å layer above and below the membrane. 0.15 M KCl was added to all systems.

We applied CGMD to observe the spontaneous diffusion of PNPLA3 to the LD which consists of a phospholipid monolayer and TAG core. CGMD simulations were performed using GROMACS 2020.5 (17) with the MARTINI 3.0 force field (18). An elastic network model was applied to maintain protein structure. The systems were minimized for 5000 steps using the steepest descent algorithm with and without a soft-core potential. Five equilibration phases followed, with lipid restraints decreasing from 200 to 10 kJ/mol/Å² and integration time steps increasing from 2 to 20 fs. During production simulations, no restraints were applied, and a 20 fs time step was used. The temperature was held at 303 K with a velocity rescaling thermostat, and surface-tension pressure coupling was implemented using a Berendsen barostat with a 4 ps coupling constant. The dielectric constant was set to 15, and long-range electrostatic interactions were truncated at 1.1 nm. Each simulation ran for 20 µs and was replicated three times.

Next, we converted the CGMD results to an AA model using the Multiscale Simulation Tool (19) and conducted all MD simulations with the Amber20 package (20). The protein was parameterized with the Amber ff14SB force field (21), POPC and DOPE lipids with Lipid14 force field (22), and TAG and SAPI lipids with the General Amber Force Field (23). The systems were minimized using the steepest descent method for 5,000 steps followed by the conjugate gradient method for 5,000 steps. Heating was performed in two stages: from 0 to 100 K over 10 ps, then to 303 K over 200 ps, with fixed lipids. The system was then equilibrated without restraints through ten 1 ns simulations at 303 K. After equilibration, 50 ns conventional MD production simulations were performed. GaMD simulations (24) included a 72 ns preparation phase with updated boost potential, followed by a 1000 ns production run with fixed boost potential. The threshold energy of GaMD was set to the lower bound, and system potential energies were averaged and recalculated every 500 ps. The boost potential was added to both the dihedral energy and the total potential energy. The upper limit of the standard deviation for the total potential energy and the dihedral energy was set to 6.0 kcal/mol. All GaMD simulations used the NPT ensemble, with pressure maintained by an anisotropic Berendsen barostat and temperature regulated by a Langevin thermostat (collision frequency of 5 ps⁻¹). The SHAKE algorithm (25) was used to restrain bonds involving hydrogen, and long-range electrostatics were handled with the particle mesh Ewald method (26) with a 12 Å cutoff. The time step of the simulation was set to 2 fs. Trajectories were collected every 10 ps for analysis.

### Fluorescence Cross-Correlation Spectroscopy (FCCS)

FCCS was performed similar to as described previously (27). Cell were transfected as above with CFP tagged hABHD5 and YFP tagged hPNPLA3 and loaded with oleic acid overnight. In brief, excitation of less than 6 μW per channel was provided by a super-continuum fiber laser (SC-Pro, YSL photonics) that was filtered into 513 nm and 561 nm channels (BrightLine FF01-513/13, FF01-561/14, Semrock), expanded, reflected by a three-band dichroic mirror (ZT442/514/561rpc-UF1, Semrock), and focused on to a diffraction-limited spot via an inverted microscope (IX83, Olympus) with a 100x oil-immersion objective (UApoN, Olympus). The excitation was focused into the center of the cytoplasm for >7 scans per cell that were >2 μm apart. The emission was filtered (FF01-485/537/627, Semrock), focused on a 40-μm diameter confocal pinhole (P40D, Thorlabs), chromatically spread by a prism (PS812-A, Thorlabs), and collected by an EMCCD camera (iXon Ultra 897, Andor). Each scan consisted 25k frames collected at 769 Hz with a 4×496 pixel region of interest, 250 EM gain, and 1 ms exposure time. Each frame of each scan was fit to the sum of the expected fluorophore emission spectra to extract the intensities versus time (I(t)) for each fluorophore.(27) The intensities versus time were correlated with the lag time (τ) according to

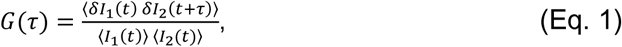

where <> represents the time average and δ represents the deviation from the scan mean. When G was calculated with a single intensity versus time (i.e., I_1_ = I_2_), G represented an autocorrelation, and when two different intensities versus time were input (i.e., I_1_ ≠ I_2_), G represented a cross-correlation. The correlations were fit to the expected shape for 2D Brownian diffusion assuming the diffusers were membrane-bound, where τ_D_ represents the dwell time, and G_0_ represents the correlation amplitude according to

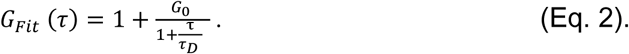

G_0_ from autocorrelations is inversely proportional to the density of diffusers within the observation spot, and the fraction of the diffusers that represent multi-colored clusters of ABHD5 and PNPLA3 (F_c_) were calculated according to

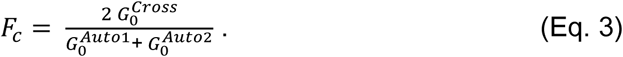

Noise within each scan and from the correlation fitting resulted in Fc values observed beyond the expected range of 0 to 1. Despite the wide range of Fc values observed for individual scans, the mean values of Fc from each condition were reasonable. Results for each condition include the combined analysis of ≥189 scans over ≥15 individual cells in ≥4 different samples on ≥4 different days. Control experiments include the analysis of AHBD5-mCherry co-expressed with YFP, WT PNPLA3-YFP co-expressed with mCherry, and PNPLA3-I148M-YFP co-expressed with mCherry, for which no cross-correlation was observed. P-values were calculated with a two-tail, unequal variance T-test.

### Statistical Analysis

All data are reported as mean ± SD. Statistical significance was determine using GraphPad Prism software version 10. Data was analyzed with Two-way ANOVA, followed by Sidak’s multiple comparisons, One-way ANOVA, followed by Sidak’s multiple comparisons, or the non-parametric Brown-Forsythe and Welch ANOVA with Dunnett’s T3 multiple comparisons for unequal variances as indicated.

## RESULTS

### Lipid droplet targeting of PNPLA3 I148M is required for the interaction with ABHD5

We previously demonstrated that the interaction between ABHD5 and PNPLA3 I148M localized to LDs (6). Murugesan *et al.* previously described basic residues in human PNPLA5 that were necessary for targeting to LDs (28). We identified a similar sequence in human PNPLA3 (^370^QRLV^373^) that was mutated to either neutral (QAAA) or neutral and acidic (AAEE) residues (Figure 1A) to examine if LD targeting of PNPLA3 I148M was required for its interaction with ABHD5. We first confirmed that the QRLV patch in PNPLA3 was required for LD targeting in EYFP fusion tags of PNPLA3 I148M in cells loaded with oleic acid overnight to promote LD formation. As expected, EYFP-tagged WT PNPLA3 partly targeted LDs as determined by co-staining cells for neutral lipids with LipidTOX Neutral red with additional cytosolic and ER localization (Figure 1B). The localization to LDs was more apparent for PNPLA3 I148M, which almost exclusively targeted LDs (Figure 1B). In contrast, PNPLA3 I148M with mutated basic residues (herein referred to as PNPLA3 I148M_QAAA_ and PNPLA3 I148M_AAEE_) were dispersed throughout the cytosol, an effect that was more apparent for PNPLA3 I148M_AAEE_ with few if any cells demonstrating localization to LDs (Figure 1B). These data indicate that PNPLA3 I148M requires basic resides for trafficking and binding to LDs.

**Figure 1:**
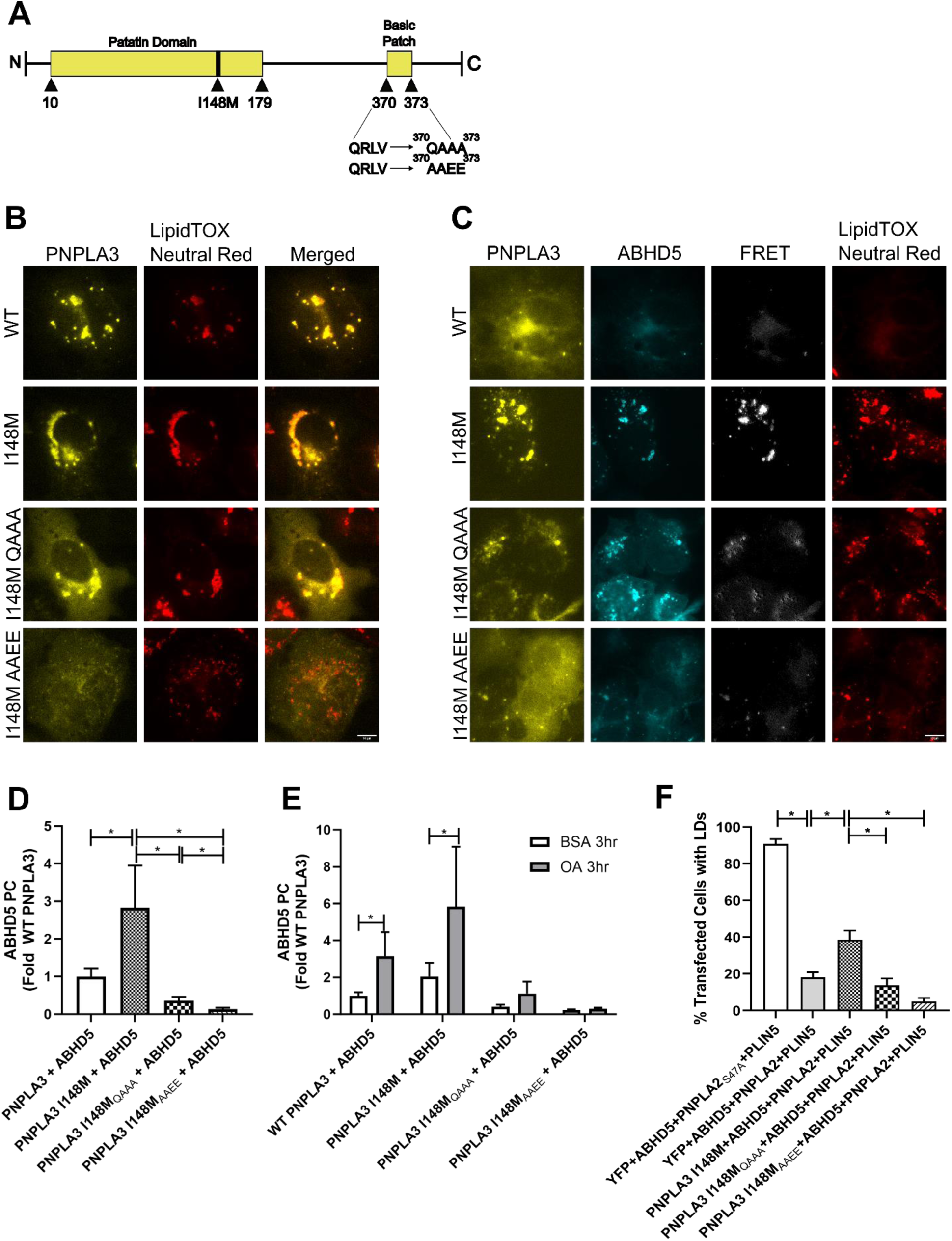
Mutations in C-terminus of PNPLA3 I148M prevent localization to lipid droplets, the interaction with ABHD5, and LD accumulation. (**A**) Schematic of human PNPLA3. (**B**) Fluorescent imaging of COS-7 cells transfected with EYFP-tagged PNPLA3 WT or mutants. Cells were treated with 0.2mM oleic acid overnight. Images are representative of results seen in three consecutive experiments. (**C**) Fluorescent imaging and FRET analysis shows EYFP-tagged PNPLA3 and ECFP-tagged ABHD5 localization and complementation in COS-7. Cells were treated with 0.2mM oleic acid overnight. Images are representative of results seen in three consecutive experiments. (**D**) Gluc protein complementation (PC) assay of HEK293A cells. Data shows that mutations cause a significant decrease in PNPLA3 and ABHD5 interaction compared to PNPLA3 I148M. Data represents averages of five trials with four technical replicates per trial. Statistics calculated using Brown-Forsythe and Welch ANOVA test with Dunnett’s T3 multiple comparisons, *p<0.05. (**E**) Gluc protein complementation (PC) assay of HEK293A cells. Data shows that a 3 hour 0.2mM oleic acid treatment significantly increases PNPLA3 and ABHD5 interaction in both WT and I148M, but causes no significant changes in the mutants. Data represents averages of three trials with four technical replicates per trial. Statistics calculated using Two-way ANOVA with Sidak’s multiple comparisons, *p<0.05. (**F**) Graph showing blinded quantification of transfected cells with visible lipid droplets after fluorescent imaging of COS-7 cells transfected with EYFP-tagged PNPLA3 I148M or mutants, mCherry-tagged ABHD5, ECFP-tagged PNPLA2 WT or ATGL_S47A_, and PLIN5. Data represents the averages of three trials. Statistics calculated using a One-way ANOVA with Sidak’s multiple comparisons, *p<0.05. Scale bar, 10 µm.

We next aimed to determine if LD targeting of PNPLA3 was required for its interaction with ABHD5. To accomplish this, we performed FRET analysis in COS7 cells co-transfected with ECFP-tagged ABHD5 and EYFP-tagged PNPLA3. As previously described (6), ABHD5 and WT PNPLA3 FRET signal was mostly on ER membranes and the FRET signal with PNPLA3 I148M was much greater and localized to neutral lipids (Figure 1C). Mutating the LD binding domain of PNPLA3 I148M to the neutral amino acids (aa; QAAA) resulted in lowered FRET signal compared to PNPLA3 I148M, levels similar to those seen in PNPLA3 WT, while the interaction of ABHD5 with PNPLA3 I148M_AAEE_ was almost completely abolished (Figure 1C). One possibility for the reduced FRET of ABHD5 with PNPLA3 I148M_QAAA_ and PNPLA3 I148M_AAEE_ could be due to the co-activator function of ABHD5 preventing LD formation and proper targeting of the QAAA or AAEE mutants. To eliminate this possibility, we expressed a mutant of ABHD5 (R299N) that cannot co-activate PNPLA2 (29). Under these conditions, the expression of PNPLA3 I148M with mouse ABHD5 R299N resulted in FRET that was localized to LDs, an effect that was reduced with PNPLA3 I148M_QAAA_, and completely abolished with PNPLA3 I148M_AAEE_ (Supplementary Figure 1A). These findings were further confirmed with Gluc Protein Complementation Assay (PCA) that allow for the monitoring of protein interactions through the reformation of luciferase activity (11). ABHD5 interacted with PNPLA3 via Gluc PC, and this effect was greater with PNPLA3 I148M. Importantly, the interaction of ABHD5 with PNPLA3 I148M_QAAA_ was significantly reduced compared to PNPLA3 I148M and almost completely abolished for PNPLA3 I148M_AAEE_ (Figure 1D). Similar protein expression levels were observed for GlucN tagged PNPLA3 and mutants (Supplementary Figure 1B). We previously demonstrated that the generation of long-chain acyl-CoAs (LC acyl-CoAs) by oleic acid supplementation increased the interaction between ABHD5 and PNPLA3 (6). The addition of oleic acid for 3 hours increased ABHD5 complementation with WT PNPLA3 and PNPLA3 I148M; however, this effect was abolished in the QRLV mutated proteins (Figure 1E), suggesting that sensing of LC acyl-CoAs required localization to membranes. Overall, these data suggest that PNPLA3 requires targeting to LDs to interact with ABHD5 and that ABHD5 itself is not sufficient to recruit PNPLA3 to LDs.

Our previous work demonstrated that PNPLA3 I148M sequesters ABHD5 to prevent co-activation of PNPLA2, the rate limiting enzyme for TAG hydrolysis (6). To investigate if PNPLA3 I148M requires LD-targeting to promote TAG accumulation, we examined LD formation in COS7 cells with overnight oleic acid loading that were transfected with mCherry-tagged ABHD5, ECFP-tagged PNPLA2, YFP-tagged PNPLA3 I148M, and PLIN5. The following day cells were monitored in a blind manner for the number of LD positive cells with brightfield microscopy. In cells transfected with PNPLA2 that lacked the active serine lipase motif (S47A), greater than 90% of cells contained LDs (Figure 1F). Expression of WT PNPLA2 in the presence of ABHD5 significantly reduced the number of cells with LDs to less than 20%, consistent with previous results (30, 31). The expression of YFP-tagged PNPLA3 I148M in the presence of ABHD5, PNPLA2 and PLIN5 increased the number of cells that contained LD compared to cells expressing ABHD5, PNPLA2 and PLIN5 alone. In contrast, the expression PNPLA3 I148M_QAAA_, or PNPLA3 I148M_AAEE_ failed to increase the number of cells that contained LD compared to cells expressing ABHD5, PNPLA2 and PLIN5. Moreover, expression of I148M_QAAA_ or I148M_AAEE_ reduced the number of cells containing LDs compared to PNPLA3 I148M (Figure 1F and Supplementary Figure 1C). Overall, these data suggest that the targeting of PNPLA3 I148M to LD is required to promote TAG accumulation in cells.

### ABHD5 requires membrane/lipid droplet targeting to interact with PNPLA3

We next asked if ABHD5 required targeting to LDs to interact with PNPLA3. We previously described a highly conserved arginine residue (R116) on the surface of ABHD5 that was necessary for binding to LDs and ER membranes (32). To determine if membrane binding of ABHD5 is required to interact with PNPLA3, COS7 cells were co-transfected with either WT mPNPLA3 or mPNPLA3 I148M and WT mABHD5 or mABHD5 R116N. WT mABHD5 interacted with WT mPNPLA3 (Figure 2A) and the FRET intensity was greatly reduced with mABHD5 R116N (Figure 2A), indicating that ABHD5 must be bound to a membrane to properly interact with mPNPLA3. mABHD5 interacted more with mPNPLA3 I148M compared to WT PNPLA3 and the FRET signal was greatly reduced with mABHD5 R116N. G0S2 was used as a negative control since it does not interact with ABHD5 (Figure 2A). The requirement for ABHD5 to bind LD and ER membranes in order to interact with PNPLA3 was further confirmed with luciferase complementation assays as the interaction of mABHD5 with both WT mPNPLA3 and mPNPLA3 I148M was greatly reduced with the R116N mutant of mABHD5 (Figure 2B). mPNPLA3-GlucN protein was expressed at similar levels (Supplementary Figure 2).

**Figure 2:**
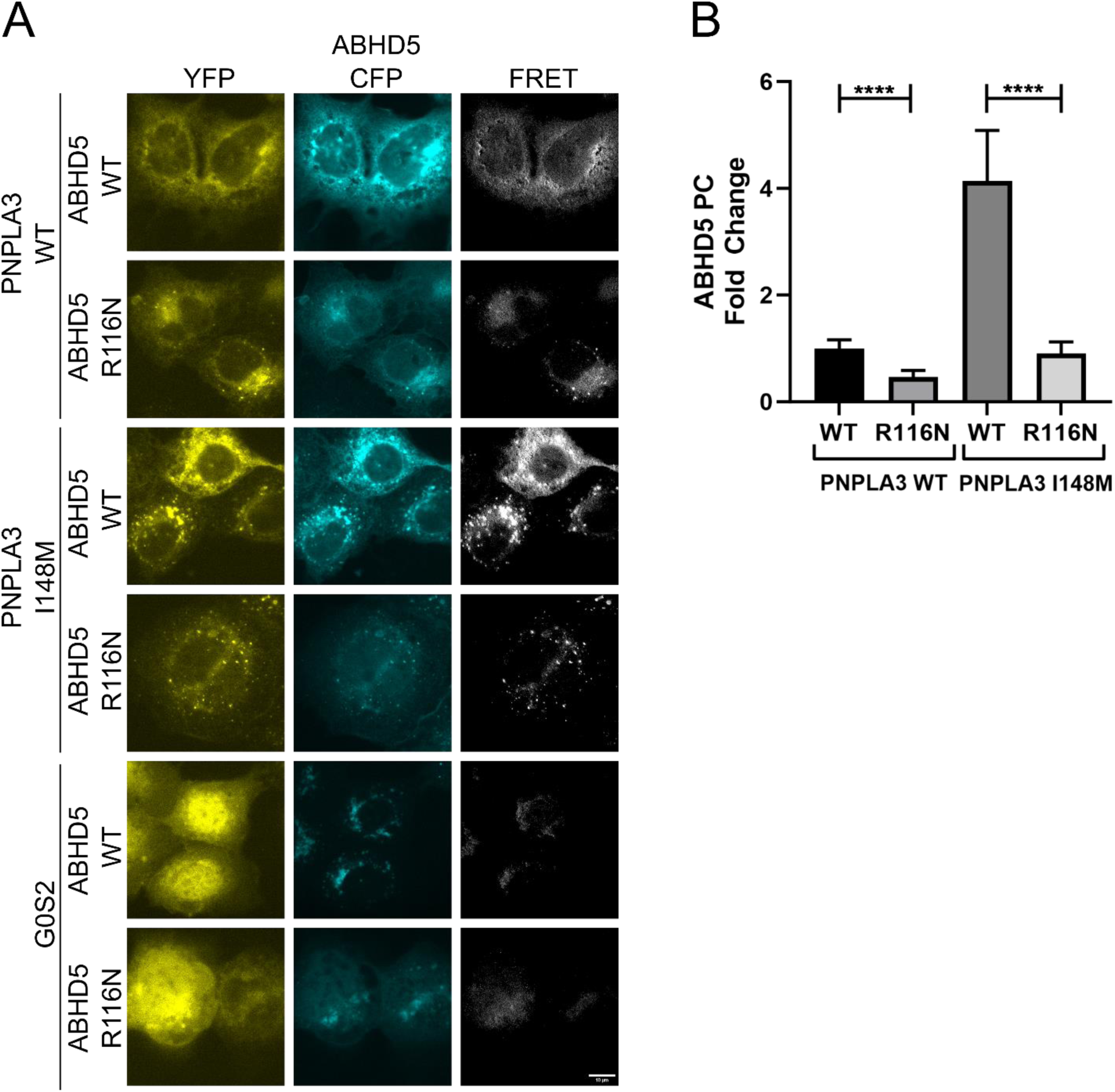
Lipid droplet/membrane targeting of ABHD5 is required to facilitate PNPLA3-ABHD5 interactions. (**A**) Fluorescent imaging and FRET analysis of COS-7 cells transfected with YFP-tagged mPNPLA3 (WT or I148M variant) or YFP-tagged G0S2, and ECFP-tagged mABHD5 (WT or R116N variant). Images are representative of results seen in four consecutive experiments. Scale bar, 10 µm. (**B**) Gluc protein complementation assay of HEK293A cells transfected with GlucN-tagged mPNPLA3 WT or mPNPLA3 I148M and GlucC-tagged mABHD5 WT or mABHD5 R116N. Data shows that cytosolic ABHD5 significantly reduces PNPLA3 and ABHD5 complementation. Data represents averages of three trials with four technical replicates per trial. Statistics calculated using Brown-Forsythe and Welch ANOVA test with Dunnett’s T3 multiple comparisons, ****p<0.001.

### Molecular modeling of PNPLA3 interaction with lipid droplets

To better understand the molecular basis for PNPLA3 LD binding we used coarse-grained molecular dynamics (CGMD) and Gaussian accelerated molecular dynamics (GaMD) to characterize the atomistic interactions between PNPLA3 and the LD. CGMD allows simulation of larger systems over longer timescales by grouping atoms into "coarse-grained" particles, while GaMD accelerates the exploration of the conformational space of a biomolecular system to reveal atomic-level interactions in greater detail. In our approach, we initially positioned PNPLA3 ∼75 Å away from the LD and used CGMD simulation to observe PNPLA3 spontaneously diffuse toward and bind to the LD. Once bound, we converted the CG to an AA model and conducted GaMD simulations to investigate the detailed interactions between PNPLA3 and the LD. Our CGMD results show that the C-terminal region of PNPLA3 (aa 345-481) is the primary area that binds to the LD (Figure 3A and 3B), consistent with our mutational analysis. In addition, three other regions in the patatin domain (aa 80-88, 147-155, and 200-225) significantly contribute to binding with the LD (Figure 3A and 3B). The first two regions are loops, while the third region consists of a helix with a loop. These regions are crucial because they may be involved in substrate binding to PNPLA3.

**Figure 3:**
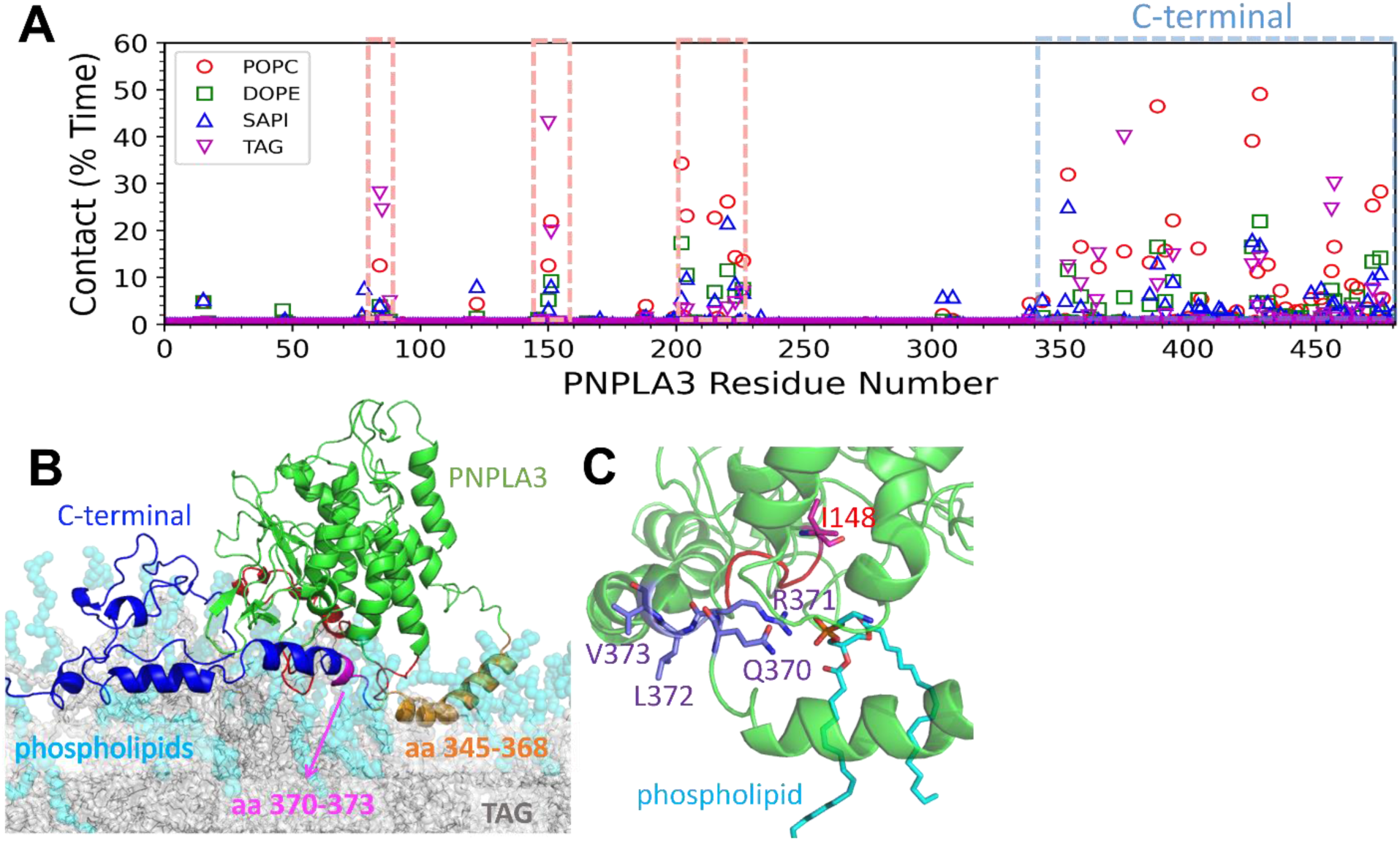
Modeling Interactions Between PNPLA3 and LDs. (**A**) The contact fraction between each PNPLA3 residue and various lipid components (POPC, DOPE, SAPI, and TAG). (**B**) The conformation of PNPLA3 bound to the LD, with PNPLA3, phospholipids, and TAG depicted in green, cyan, and gray, respectively. The three regions (aa 80-88, 147-155, and 200-225), which form interactions with the LD, are highlighted in red. The C-terminal of aa 369-428 and 433-481 is shown in blue. The helix of aa 345-368 and aa 370-373 is shown in orange and magenta, respectively. (**C**) GaMD snapshot of WT PNPLA3 bound to the LD, I148, the ^370^QRLV^373^ motif, and the phospholipid interacting with R371, colored in magenta, purple, and cyan, respectively.

We conducted GaMD simulations to explore the atomistic details of protein-membrane interactions. Our results revealed that four amphipathic helices (aa 345-368, 369-376, 377-387, and 388-402) of the C-terminus of PNPLA3 play a crucial role in targeting the LD. Notably, the helix formed by aa 345-368 of PNPLA3 is particularly significant, as it is fully buried within the membrane and interacts with TAG (Figure 3B). Upon PNPLA3 binding to the LD, the regions formed by aa 80-88 and 147-155 of the patatin domain interact with TAG and phospholipids. Moreover, the region formed by aa 200-225, including the helix from aa 215-223, also interacts mostly with phospholipids, but also TAG. Comparing the conformations of these regions before and after LD binding, we found that all regions interacting with the LD, especially the C-terminus of PNPLA3 undergo significant conformational changes upon binding (Figure 3). Once bound, the LD restricts PNPLA3 to a specific conformation, which may facilitate ABHD5 recognition of PNPLA3. This rigidity may also contribute to the stabilization of the ABHD5-PNPLA3 complex on the LD surface. Our modeling results highlight the specific regions of PNPLA3 that form direct interactions with the LD, aligning well with experimental findings that the C-terminal region is pivotal for LD targeting.

GAMD simulations identified that the ^370^QRLV^373^ motif plays a critical role in PNPLA3-LD interactions. This motif embeds deeply into the phospholipid layer, where R371 forms polar interactions with the phospholipid heads (Figure 3C), while L372 and V373 engage in hydrophobic interactions with the phospholipid tails. The structural arrangement of ^370^QRLV^373^ explains why mutating to neutral (QAAA) or acidic (AAEE) experimentally reduces PNPLA3 binding to LDs. Overall, molecular simulations provides insights into how PNPLA3 binds LD and atomistic detail of the QRLV in interacting with phospholipids.

We previously noted a stronger interaction between PNPLA3 I148M with ABHD5 compared to WT PNPLA3 (6) ( see also Figure 1 and 2) which could be due to greater binding between ABHD5 and PNPLA3 I148M or due to greater expression of PNPLA I148M (33). To understand the molecular basis for the greater interaction we performed live cell dual color fluorescent cross-correlation spectroscopy (FCCS) to determine the fluorescent correlation (F_c_) and diffusion rates of the interacting proteins. In transfected cells, FCCS analysis demonstrated greater cross-correlation between ABHD5 and PNPLA3 I148M compared to WT PNPLA3 (F_c_ of 0.32 ± 0.04 and 0.57 ± 0.04 for the WT and variant, respectively (p < 0.0001); Figure 4A and B and C). Further inspection demonstrates that 78 ± 7 % and 83 ± 6 % of the WT PNPLA3 and PNPLA3 I148M, respectively, were co-diffusing with ABHD5 with no significant difference between them. The difference in the F_c_ is dominated by the fraction of the ABHD5 that was co-diffusing with the PNPLA3, which was 24 ± 3 % and 45 ± 5 % for the WT and I148M, respectively (p = 0.0005; Supplementary Figure 3). Moreover, the diffusion of PNPLA3 I148M and ABHD5 complexes was slower compared to WT PNPLA3 and ABHD5 complexes (Figure 4D). Interestingly, FCCS indicates that there are fewer independent diffusers of PNPLA3 I148M than WT PNPLA3 despite the greater expression levels of I148M than the WT suggesting that multiple copies of the variant may be co-diffusing with ABHD5. These data are consistent with the observed slower diffusion rate of the multi-color complexes with the I148M variant diffuse compared to those with WT. Overall, single-molecule analysis suggests a stronger association between ABHD5 and PNPLA3 I148M than ABHD5 and WT PNPLA3.

**Figure 4:**
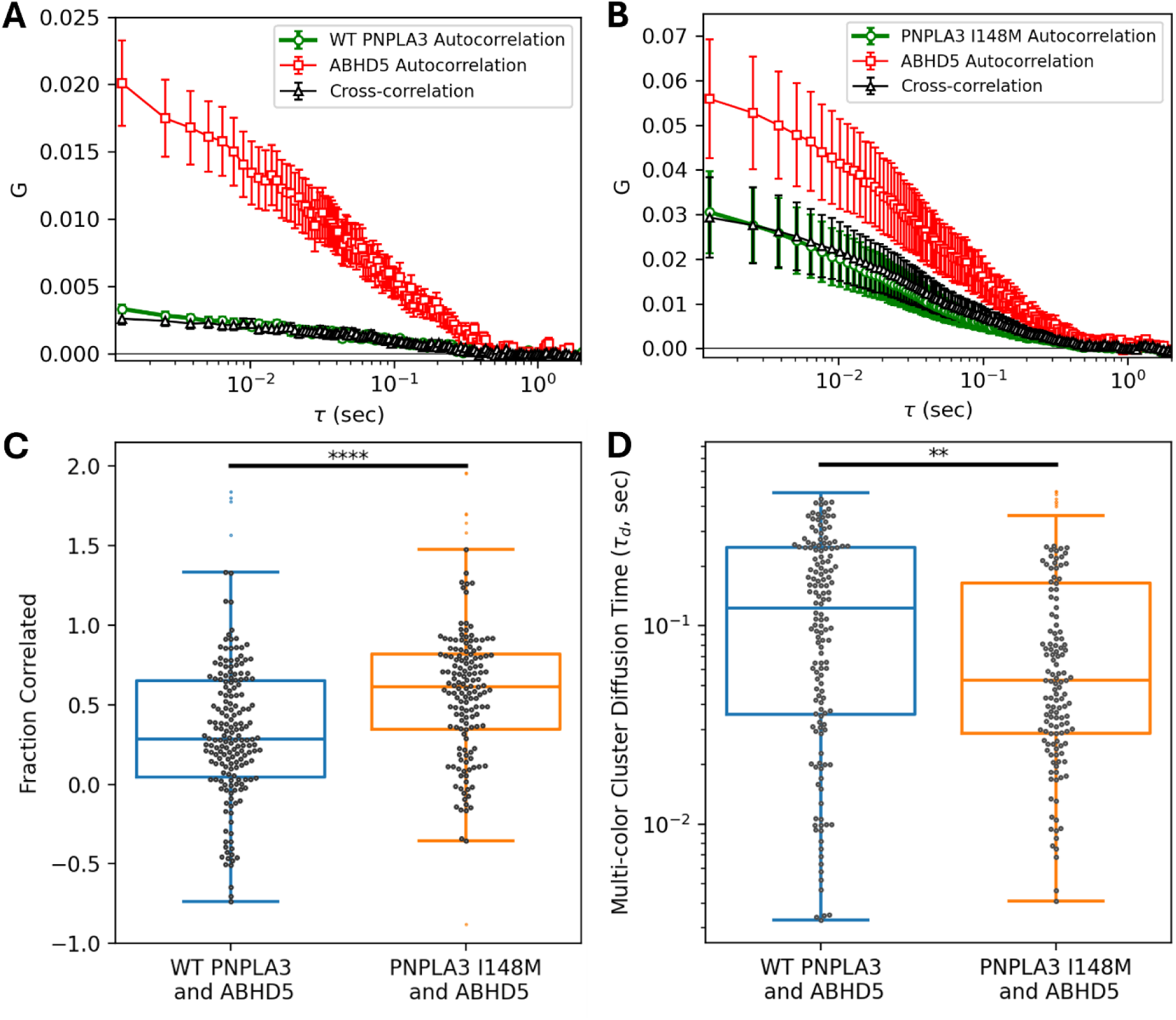
Fluorescence Cross Correlation Spectroscopy of ABHD5 with WT PNPLA3 and PNPLA3 I148M. FCCS was performed to analyze the co-diffusion of ABHD5 and PNPLA3. The mean and standard error of the mean of the autocorrelations and cross-correlations for (**A**) ABHD5 and WT PNPLA3 and (**B**) ABHD5 and PNPLA3 I148M are shown. Larger amplitude correlations, greater similarity between correlation amplitudes, and faster cluster diffusion in the presence of the variant versus WT are significant. (**C**) A greater fraction of ABHD5 and the variant were co-diffusing than ABHD5 and WT PNPLA3 (p = 0.00002). (**D**) The multi-color clusters of ABHD5 and PNPLA3 demonstrated a slower diffusion time with WT PNPLA3 than the variant (p = 0.002). (**C-D**) Noise within each collected scan and uncertainties of the fits resulted in a wide distribution of individual scan fit results (*symbols*). Boxplots show the median and extent of the data (*lines*).

### The lipid droplet binding domain of PNPLA3 partly re-constitutes the interaction of PNPLA4 with ABHD5

PNPLA enzymes belong to a large family of lipases and are marked by their highly conserved patatin domain that contains a catalytic dyad (34). PNPLA4 is ubiquitously expressed and has been shown to possess triglyceride lipase and transacylase activity (35). However, PNPLA4 lacks the C-terminus of many of the other PNPLA enzymes and localizes to mitochondria (36) where it is thought to function as a retinol transacylase (37, 38). We previously demonstrated that ABHD5 interacted with PNPLA3 but not with PNPLA4 (6). We investigated if subcellular localization to LDs determines the ability to interact with ABHD5 by adding the C-terminus which contains the LD-targeting domain of PNPLA3 to PNPLA4 which we hypothesized would be a gain of function for the interaction with ABHD5. We therefore fused the C-terminus of PNPLA3 onto the end of PNPLA4 (PNPLA4-PNPLA3) as well as the C-terminus with mutated basic residue (PNPLA4-PNPLA3_AAEE_) (Figure 5A). We then investigated the subcellular targeting of PNPLA4 along with the fusion proteins in COS7 cells loaded with oleic acid overnight. As expected, WT PNPLA3 targeted LDs and PNPLA4 was localized mostly to mitochondria as determined by staining with LipiBlue and Mitotracker Red, respectively (Figure 5B). Fusion of the C-terminus of PNPLA3 to PNPLA4 (PNPLA4-PNPLA3) promoted LD targeting (Figure 5B).

**Figure 5:**
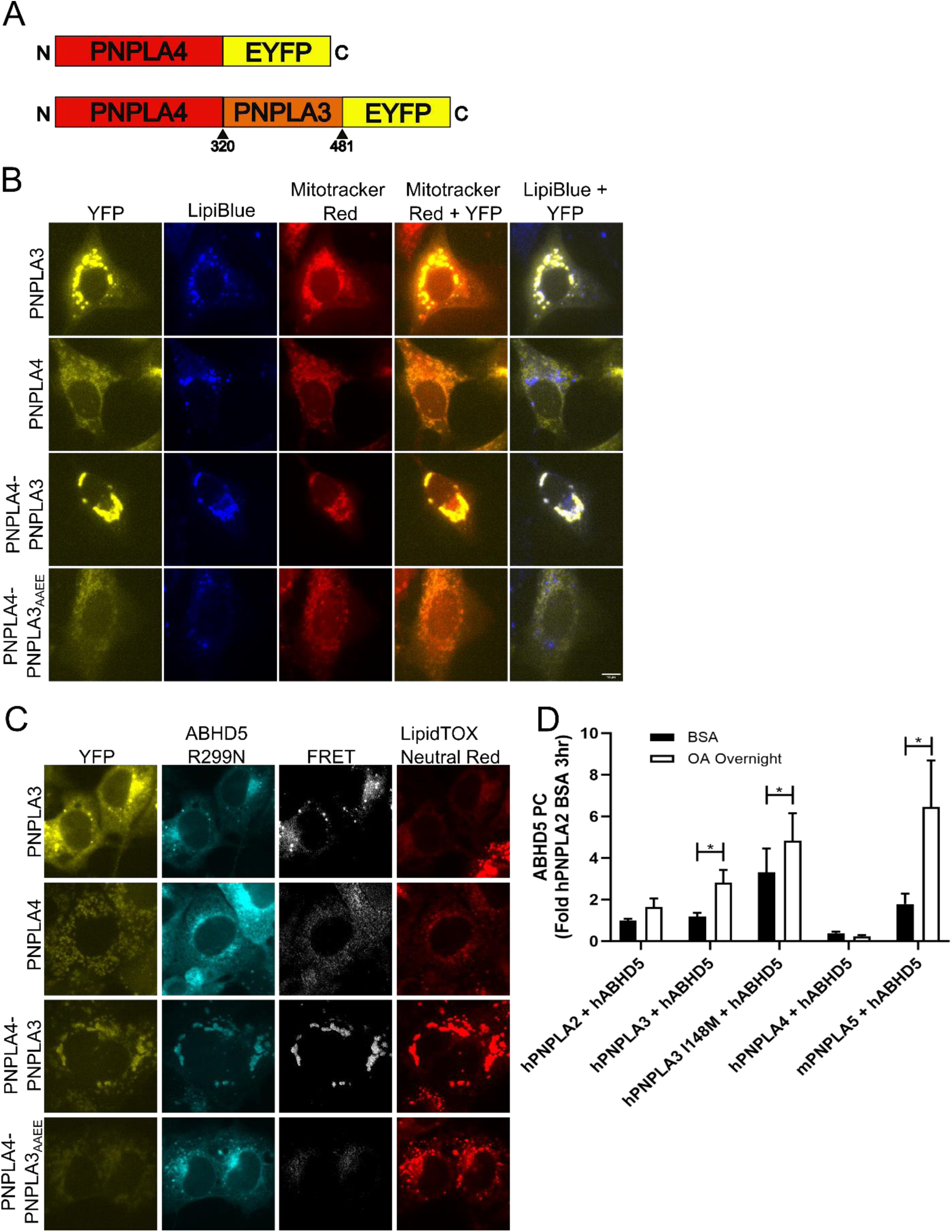
ABHD5 interacts with multiple PNPLA family members. (**A**) Schematic of PNPLA4 fusions. (**B**) Fluorescent imaging of U2OS cells transfected with YFP-tagged PNPLA4, PNPLA3 WT, PNPLA3-PNPLA4 fusion, or PNPLA3-PNPLA4 fusion_AAEE_ mutant to show lipid droplet targeting. Cells were treated with 0.2mM oleic acid overnight and stained with Mitotracker Red and LipiBlue. Images are representative of results seen in three technical replicates of three experiments. (**C**) Fluorescent imaging and FRET analysis of COS-7 Cells transfected with YFP-tagged PNPLA4, PNPLA3 WT or PNPLA3 mutant and ECFP-tagged ABHD5 R229N. Cells were treated with 0.2mM oleic acid overnight and stained with LipidTOX Neutral Red. Images are representative of results seen in three technical replicates of three consecutive experiments. (**D**) Gluc protein complementation assay of HEK293A cells transfected with various PNPLA family proteins tagged with GlucN and ABHD5-GlucC. Cells were treated with 0.2mM oleic acid in 10% BSA or 10% BSA alone overnight. Data shows that oleic acid treatment significantly alters the interaction of ABHD5 with PNPLA3 WT, PNPLA3 I148M and PNPLA5. Data represents averages of three trials with four technical replicates per trial. Statistics calculated using Two-way ANOVA with Sidak’s multiple comparison test, *p<0.05. Scale bar, 10 µm.

Mutating the basic residues to acidic in PNPLA3 (PNPLA4-PNPLA3_AAEE_) eliminated LD targeting and resulted in localization mostly to mitochondria (Figure 5B).

Next, we examined if targeting of PNPLA4 to LDs led to a gain of function for ABHD5 binding. The EYFP-tagged proteins were co-transfected with ECFP-tagged ABHD5 R299N which prevents activation of PNPLA2, thereby promoting LD formation. FRET analysis demonstrated the interaction between PNPLA3 and ABHD5, but very little to no FRET with PNPLA4 (Figure 5C). Greater FRET intensity was observed with the PNPLA4-PNPLA3 fusion compared to those transfected with PNPLA4, and the effect was abolished in the PNPLA4-PNPLA3_AAEE_ fusion. These data indicate that the LD/ER targeting is needed for ABHD5 interaction with PNPLA family members.

### ABHD5 interacts with additional PNPLA family members

As mentioned above, the PNPLA family of proteins consists of 9 enzymes, all containing the patatin-like protein structure, which houses the catalytic domain of these serine hydrolases (39). ABHD5 co-activates PNPLA2 (ATGL) (40) interacts with PNPLA3 (6) and the functional interaction between ABHD5 and PNPLA1 is important for skin barrier formation (41, 42).

Therefore we aimed to determine if other members of the PNPLA family associate with hABHD5 in luciferase PCA. Cells were treated with either BSA (control) or oleic acid complexed with BSA overnight to mimic nutritional feeding and LD formation (Figure 5D). As reported previously, hPNPLA2 interacted with hABHD5 and this effect was not modified by oleic acid (6). Both WT hPNPLA3 and hPNPLA3 I148M interacted with ABHD5 and this effect was increased with oleic acid supplementation (Figure 5D). Consistent with previous data (6), hPNPLA4 did not interact with ABHD5. mPNPLA5, the closest related family member to PNPLA3 which rose from a gene duplication in mammals (43), also interacted with ABHD5 and showed a significant increase with oleic acid. Human PNPLA8, did not express properly (Supplementary Figure 4A), and we were not able to evaluate the interaction with ABHD5. Human PNPLA6, a phospholipase with neuronal function (44), human PNPLA7, a phospholipase with hepatic function (45, 46), and human PNPLA9 (47) while expressed properly (Supplementary Figure 4A); did not demonstrate a detectable interaction with ABHD5 in this assay (Supplementary Figure 4B).

### LD targeting of PNPLA3 I148M is required to promote liver steatosis

Previous work demonstrated that the I148M mutation is not a simple loss of function mutation, but a gain of function as knock-out of PNPLA3 does not produce steatosis in mice (48). Overexpression of hPNPLA3 I148M in the liver or transgenic mice with the PNPLA3 I148M substitution develop liver steatosis (49, 50). To determine if LD targeting of hPNPLA3 I148M is required for the liver steatosis, we utilized PNPLA3-KO mice and re-expressed WT hPNPLA3, hPNPLA3 I148M, I148M_AAEE_ or GFP control specifically in the liver of mice and challenged them with a diet high in sucrose to induce liver steatosis (49). As expected, the expression of hPNPLA3 I148M increased liver TAG levels compared to mice expressing GFP and WT hPNPLA3, and the expression of WT hPNPLA3 had no effect on liver TAG levels compared to GFP (Figure 6A). The expression of hPNPLA3 I148M_AAEE_ resulted in lower TAG levels compared to hPNPLA3 I148M and was not significantly different compared to mice with GFP expression (Figure 6A). The liver steatosis was confirmed with oil red O staining of frozen liver sections which demonstrated greater steatosis in mice with expression of hPNPLA3 I148M (Figure 6B). We confirmed the cellular localization of the hPNPLA3 proteins in fractionated liver. No hPNPLA3 signal was detected in GFP mice and much of the WT hPNPLA3 and hPNPLA3 I148M localized to the LD fraction. In contrast, no hPNPLA3 I148M_AAEE_ was found in the LD fraction and the majority of the signal was found in cytosol (Figure 6C). PLIN2 served as markers for LD, Mek1/2 for cytosol and AIF for the Mitochondria (Mito) and membranes (MB) fraction (Figure 6C). Overall, these data suggest that LD targeting of PNPLA3 I148M and the interaction with ABHD5 is required to promote liver steatosis.

**Figure 6:**
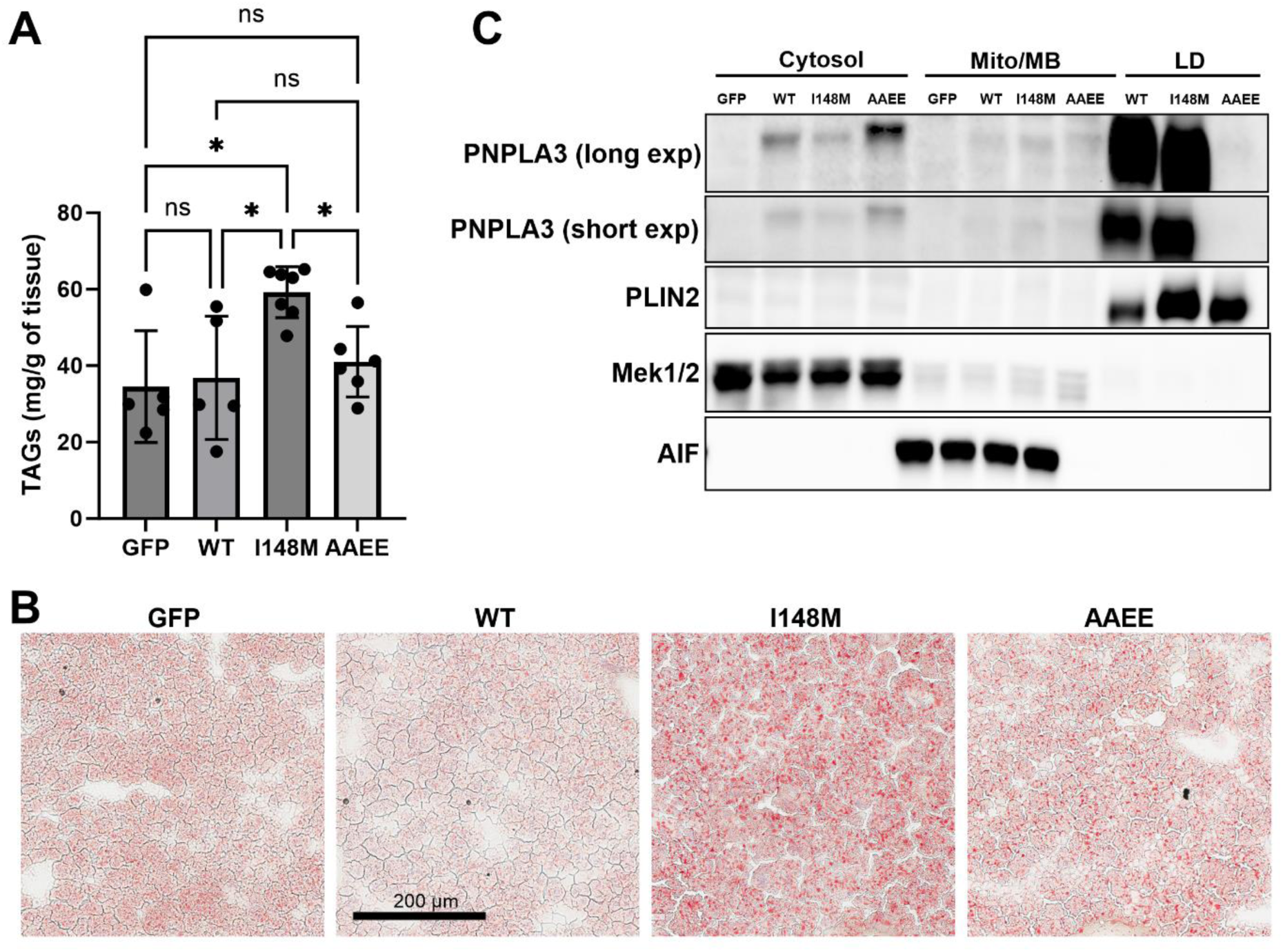
PNPLA3 I148M targeting to LDs is required to promote liver steatosis. (**A**) Liver TAGs were quantified from PNPLA3-KO mice with liver expression of GFP, WT PNPLA3 (WT), PNPLA3 I148M (I148M), or PNPLA3 I148M_AAEE_ (AAEE). (**B**) Representative Oil Red O staining of frozen liver sections from mice in A. (**C**) Livers from mice in A were fractionated into cytosol, mitochondria/membrane (Mito/MB), and lipid droplets (LD). Fractionation data are representative from three independent experiments. *p<0.05, **p<0.01 as determined by One-way ANOVA with Sidak’s multiple comparison test. Scale bar, 200 µm.

## DISCUSSION

PNPLA2, the closely related family member of PNPLA3, is initially synthesized in ER membranes and transferred to LDs, classifying it as an ERTOLD protein that contains basic residues that makeup hydrophobic hairpin structures that embed in bilayer membranes (51). We found that mutating the basic residues in the C-terminus of PNPLA3 I148M prevented its localization to LDs and interaction with ABHD5. In contrast, ABHD5 is initially synthesized in the cytosol and contains amphipathic sequences that enable its transient targeting to the LDs, classifying it as a “CYTOLD” protein (51). Indeed, R116 in ABHD5 stabilizes an amphipathic helix for membrane binding (52). Thus, the mechanisms by which ABHD5 and PNPLA3 target LD differs; however, both interaction on LDs. Recently Sherman et al. investigated the ER targeting of PNPLA3, demonstrating that PNPLA3 does not contain an ER-transmembrane domain; however, additional membrane targeting mechanisms, such as the role of a basic residue, were not investigated (53). Interestingly, endogenously tagged PNPLA3 was primarily found in the Golgi of Hep3B hepatocytes, while PNPLA3 I148M was localized mostly to LD upon oleic acid loading (53). Interestingly, we previously noted that EYFP-tagged PNPLA3 localized to LDs within or adjacent to Golgi (6). The physiological significance of PNPLA3 localization to Golgi is currently unclear.

In our previous and current work we noted that PNPLA3 I148M interacts more with ABHD5 than WT PNPLA3 which could be due to greater binding of ABHD5 for PNPLA3 I148M or greater protein expression and stability (6, 33). Our FCCS analysis confirms the greater association between ABHD5 and PNPLA3 I148M than ABHD5 and WT PNPLA3 even when the varying expression levels are considered (Figure 4). The increased interaction between ABHD5 and PNPLA3 I148M compared to WT PNPLA3 is not likely due to differences in protein expression as total protein expression was similar (Supplementary Figure 1C and Supplementary Figure 2). Rather, data suggest that ABHD5 and PNPLA3 I148M form a more stable complex as noted by slower diffusion compared to WT PNPLA3. Experiments are underway to understand the atomistic basis of PNPLA3 I148M stabilization and if binding to LD may be important for binding to ABHD5.

Previous work identified a surface patch in ABHD5 in which the residue R116 was found to be essential for membrane binding. Mutating this position from an arginine to an asparagine (R116N) disrupts hydrogen bonding of nearby lysine residues and abolishes the ability of ABHD5 to target to the membranes (32, 52); however, does not affect the ability of ABHD5 to co-activate PNPLA2 or interact with PLIN1 or 5 (32). In this work, we have shown that the R116N impaired ABHD5 membrane localization and significantly reduced the interaction of ABHD5 with WT PNPLA3 and PNPLA3 I148M. Membrane binding of ABHD5 may expose ABHD5 residues that allows for greater interaction with PNPLA3, that is not required for PNPLA2 activation or PLIN interactions and could be important for differentially regulating ABHD5 partner binding and cellular localization. Further work is required to understand the structural significance of ABHD5 and PNPLA3 membrane binding and how this affects their interaction which would have important repercussions in how the I148M variant causes MASLD. ABHD5 interacts more with PNPLA3 than PNPLA2, and PNPLA3 I148M sequesters ABHD5, significantly inhibiting PNPLA2-dependent lipolysis (6). The R116N mutation does not disrupt the co-activation function of ABHD5 on PNPLA2, suggesting that the co-activation function of ABHD5 is not sufficient to promote the interaction with PNPLA3. The exact biophysics by which membrane binding of PNPLA3 promotes its interaction with ABHD5 is currently unknown. Interestingly, MD simulations and structural modeling identified a similar patch of residues that were recently identified in PNPLA3 that is required for LD binding and the association with ABHD5 (54).

We found that fusing the C-terminus of PNPLA3 including the LD binding domain to PNPLA4 effectively reconstituted the ABHD5 interaction with PNPLA4 and mutating the basic residues in LD binding domain prevented the interaction with ABHD5. Taken together, these data indicate that monolayer membrane binding is required for ABHD5 to interact with PNPLA family proteins and that the patatin domain likely constitutes the minimal domain required to interact with ABHD5. It is currently unknown if mammalian ABHD5 localizes to mitochondria, however the ancestral ABHD4/5 in drosophila localizes to mitochondria; but this function seems to have been lost during evolution. Whether mammalian ABHD5 localizes to mitochondria is currently unclear, but co-localization seems to provide a mechanism for ABHD5 to interact with PNPLAs.

ABHD5 interaction with other PNPLA family members was further investigated via luciferase PCA. Interestingly, our data indicate that ABHD5 interacts with lipases (PNPLA2, 3 and 5) but not phospholipase members of PNPLAs (PNPLA6, 7 and 9). PNPLA5 is involved in the initiation of autophagy in which LDs are used to supplement autophagosome membrane formation (55). Furthermore, PNPLA5 mRNA decreases during fasting, increases during adipocyte differentiation, and is upregulated in the liver with high sucrose and high fat feeding (56, 57). We found that ABHD5 interaction with PNPLA5, similar to PNPLA3 was increased with oleic acid supplementation that mimics the nutritionally fed state. PNPLA5 localizes to LDs and is structurally and evolutionarily similar to PNPLA3 (34) and the expression pattern of *Pnpla5* is highly correlated to that of *Pnpla3* (57). This similarity in expression and localization of PNPLA5 and PNPLA3 may indicate a similar physiological function. Together, these data suggest the ABHD5/PNPLA5 interaction may also have a role in the nutritional fed state. Previous work found ABHD5 interacted with PNPLA9 in IP, but direct interaction was not evaluated (58).

Our previous work demonstrated that PNPLA3 I148M sequesters ABHD5 away from PNPLA2, thereby preventing lipolysis activation (6). PNPLAs are thought to physically associate with LDs to carry out their function (28) and by abolishing the LD binding domain of PNPLA3 I148M this prevented the interaction with ABHD5. As a result, the number of cells with neutral lipid accumulation in LDs was significantly reduced compared to those transfected with PNPLA3 I148M with the LD binding domains intact. This suggests that PNPLA3 I148M must physically associate with LDs to sequester ABHD5 and promote TAG accumulation. Our data also suggest that targeting of PNPLA3 I148M on LD is required for promoting liver steatosis.

The I148M substitution of PNPLA3 results in a reduction in lipase activity, suggesting that the mutation renders a partial loss of function (59). However, whole body deletion of PNPLA3 does not cause steatosis, implying that I148M is not a simple loss of function. Normally the expression of PNPLA3 is regulated at the nutritional level where mRNA expression is highest under fed conditions, but lower under fasted conditions in the liver (60). However, the PNPLA3 I148M substitution renders the mutant protein resistant to proteasomal degradation (61) through the evasion of ubiquitination as a synthetic mutant of PNPLA3 that evades ubiquitination, but maintains enzymatic activity is sufficient to promote TAG accumulation (33). Thereby, the resistance of the mutant protein to proteasomal degradation allows it to escape the normal nutritional regulation and to accumulate on LDs and provides and additional explanation of how PNPLA3 I148M promotes steatosis. We mutated a basic amino acid in the C-terminus of PNPLA3 I148M, which prevented the mutant protein from targe LDs and thereby promoting hepatosteatosis. Together these data suggest that PNPLA3 I148M targeting and accumulation on LDs is required for initiating steatosis and further support a model by which PNPLA3 I148M sequesters ABHD5 and prevents activation of PNPLA2. LC-acyl-CoAs are endogenous ligands of ABHD5 (62) and function to allosterically regulate the interaction of ABHD5 with PLIN1,5 and PNPLA3. Preventing PNPLA3 targeting to LDs also blocked the ability of oleic acid supplementation to increase the interaction between ABHD5 and PNPLA3 suggesting that initial ABHD5-PNPLA3 complex formation or PNPLA3 membrane binding is required for LC-acyl-CoAs to increase the ABHD5-PNPLA3 interaction.

Johnson et al. demonstrated that PNPLA3 facilitates the release of VLDL as knockdown or knockout of liver PNPLA3, resulting in TAG accumulation under lipogenic conditions (63). Overall, these data suggest that PNPLA3 I148M is a loss of function mutation; however, this is a simplistic interpretation and the I148M substitution likely represent a neomorph (64).

Interestingly, the authors also found that overexpression of PNPLA3 I148M promoted liver TAG accumulation and that knockin mice for PNPLA3 I148M fed a western diet supplemented with glucose and fructose have reduced TAG secretion. Overall, these data suggest that while the WT enzyme has a function in VLDL secretion under lipogenic conditions, the mutant protein also functions to disrupt TAG hydrolysis. Thus, both loss of function in VLDL secretion and gain of function to sequester ABHD5 could contribute to TAG accumulation in patients with PNPLA3 I148M. Interestingly, ablation of ABHD5 in vitro or in vivo reduces liver TAG secretion (2, 65), suggesting that sequestration of ABHD5 by PNPLA3 I148M prevents its function in VLDL secretion. Current experiments are underway to investigate how PNPLA3 I148M might disrupt the function of ABHD5 in TAG secretion.

Recently, our model in which PNPLA3 I148M sequesters ABHD5 to initiate steatosis (6) was evaluated by the group of Hobbs and Cohen and provided additional evidence that overexpression of ABHD5 can reduce steatosis induced by PNPLA3 I148M. Here the authors also identified that membrane/LD binding of ABHD5 and PNPLA3 is important for the interaction, although the authors identified different patches in ABHD5 (66) and PNPLA3 that mediated LD-targeting. PNPLA3 is of therapeutic interest and a recent clinical trial demonstrates that targeting PNPLA3 in patient that are homozygous for the I148M variant with siRNA reduces steatosis (67). Understanding the biophysical basis for the ABHD5/PNPLA3 interaction could provide additional therapeutic avenues.

## DATA AVAILABILITY

All data reported in this paper will be shared by the corresponding contact.

### Author Contributions

E.P.M. conceived the experiment, E.P.M., G.T. and J.G.G. designed the experiments. G.T, N.T., A.J.B., C.V.K., A.K., Y.M.H. and E.P.M. conducted the experiments. G.T., E.P.M., A.K., C.V.K. and Y.M.H. analyzed the data and E.P.M., G.T. C.V.K. and Y.M.H, wrote and edited the manuscript. J.W.P. and J.G.G. provided critical reagents. All authors read the manuscript and approved the final version.

### Funding Information

This work was supported by grant R01 DK126743 and P30DK092926 (Michigan Diabetes Research Corridor) from the National Institute of Diabetes and Digestive and Kidney Diseases to EPM, DK076229 to JGG and CVK from the National Institute of Diabetes and Digestive and Kidney Diseases of the National Institutes of Health, and the Barber Integrative Metabolic Research Program (EPM, CVK, JGG and YHM). AJB was supported by grant 2T32HL120822 from the National Heart, Lung, and Blood Institute of the National Institutes of Health awarded to Wayne State University. The modeling work was supported by the Wayne State University Startup Fund to YMH. We thank Wayne State University High Performance Computing Center for their support of all simulations. The funding agencies were not involved in the study design, collection analysis and interpretation of data; in the writing of the report; and in the decision to submit the article for publication.

## Abbreviations

AA: all-atom
ABHD5: Alpha-beta hydrolase domain-containing 5
ATGL: Adipose triglyceride lipase
BSA: Bovine serum albumin
CGMD: coarse-grained molecular dynamics
DAG: Diacylglycerol
DOPE: Dioleoylphosphatidylethanolamine
FA: Fatty acid
FFA: Free fatty acid
FCCS: fluorescent-cross correlation spectroscopy
FRET: fluorescence resonance energy transfer
GaMD: Gaussian accelerated molecular dynamics
Gluc: *Gaussia* luciferase
LD: lipid droplet
MASLD: metabolic dysfunction–associated steatotic liver disease
PCA: protein complementation assays
PLIN1: Perilipin-1
PNPLA2: Patatin Like Phospholipase Domain Containing 2
PNPLA3: Patatin Like Phospholipase Domain Containing 3
POPC: palmitoyl-2-oleoyl-glycero-3-phosphocholine
PCA: protein complementation assays
SAPI: 1-stearoyl-2-arachidonoyl-phosphatidylinositol
TAG: Triacylglycerol

