## Supplemental Information for "Lipid droplet targeting of ABHD5 and PNPLA3 I148M is required to promote liver steatosis"

**Supplementary Figure 1**

**Supplementary Figure 2**

**Supplementary Figure 3**

**Supplementary Figure 4**

**Supplementary Table S1**

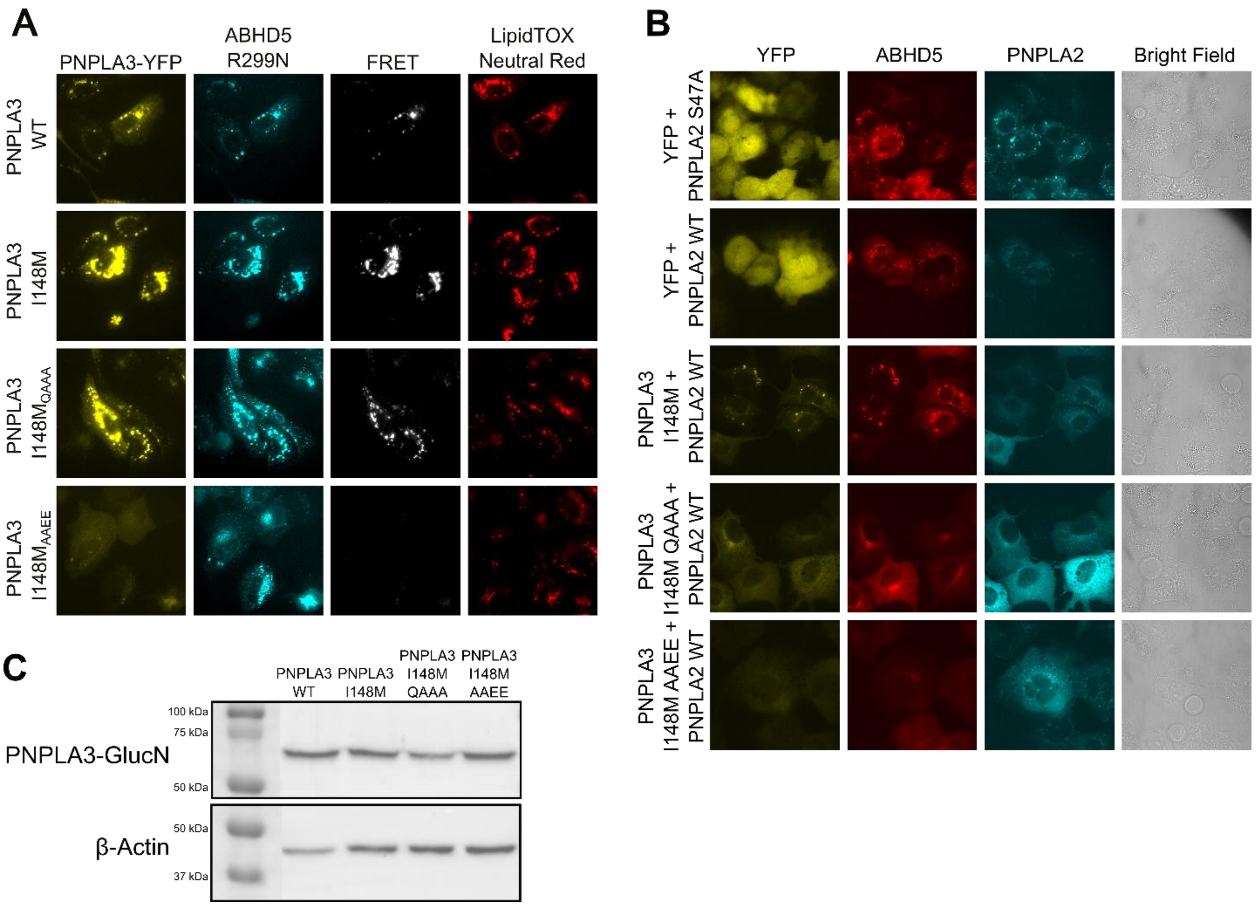


**Supplementary Figure 1:** (**A**) Fluorescent imaging and FRET analysis of U2OS cells shows localization of EYFP-tagged WT hPNPLA3 or mutants and ECFP-tagged mABHD5-R299N. Cells were treated with 0.2mM oleic acid overnight and stained with LipidTOX Neutral Red. Images are representative of three consecutive experiments with three technical replicates per trial. Scale bar, 10 µm. (**B**) Representative panel of images used for blinded quantification of transfected cells with visible lipid droplets after fluorescent imaging of COS-7 cells transfected with EYFP-tagged PNPLA3 I148M or mutants, mCherry-tagged ABHD5, ECFP-tagged PNPLA2 WT or $ATGL_{S47A}$, and PLIN5. Cells were treated with 0.2mM oleic acid overnight. Images are representative of three trials with three technical replicates per trial. Scale bar, 10 µm. (**C**) Western blot of PNPLA3-GlucN and β-Actin expression in HEK293A cells transfected with plasmids used for protein complementation assay. Blot shows similar PNPLA3-GlucN expression for variants and is representative of four separate western blot experiments.

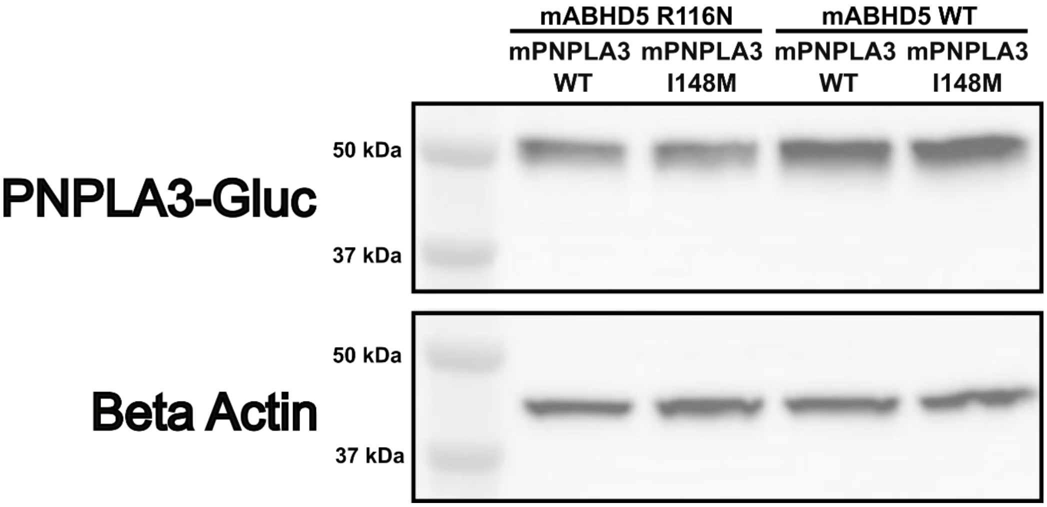


**Supplementary Figure 2:** Western blot of PNPLA3-GlucN and β-Actin expression in HEK293A cells transfected with plasmids used for protein complementation assay. Blot shows that PNPLA3-GlucN expression is present and similar across experimental treatments.

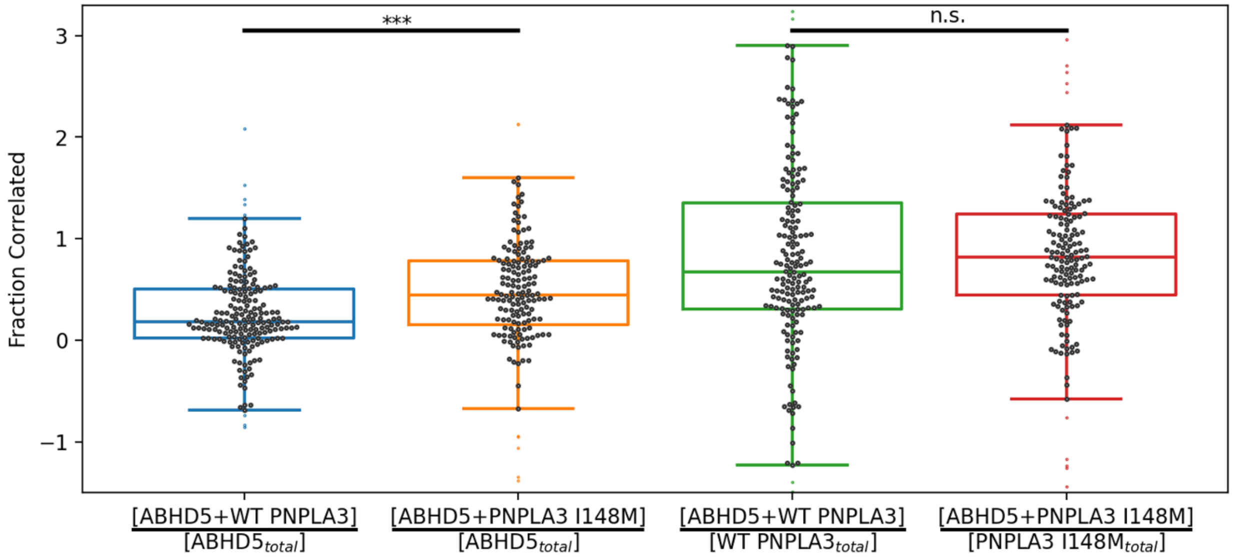


**Supplementary Figure 3:** Analysis of the correlation amplitudes revealed the fraction of each protein that is part of a multi-color protein cluster through calculating $G_0^{Cross} / G_0^{Auto}$. ABHD5 is more likely to co-diffuse with PNPLA3 I148M than with WT PNPLA3 ($p = 0.0005$). The WT and the variant PNPLA3 are indistinguishably likely to be co-diffusing with ABHD5. Noise within each collected scan and uncertainties of the fits resulted in a wide distribution of individual scan fit results (symbols). Boxplots show the median and extent of the data (lines).

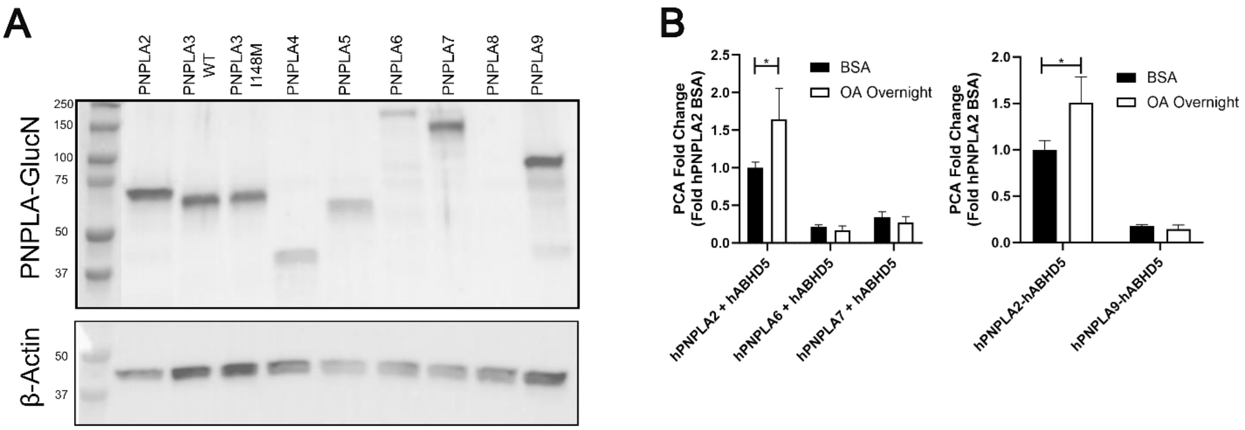


**Supplementary Figure 4:** (**A**)Western blot showing expression of PNPLA proteins with GlucN tag and β-Actin. With the exception of PNPLA8, all PNPLAs were expressed in the protein complementation assays. (**B**) Gluc PC assay of HEK293A cells showing the lack of interaction of GlucN-tagged PNPLA6, PNPLA7 and PNPLA9 with GlucC-tagged ABHD5, both at baseline and after overnight 0.2mM oleic acid treatment. Data represents the average of 3 trials with 4 technical replicates per trial. Statistics calculated using Two-way ANOVA with Sidak’s multiple comparison test (*p<0.05).

**Supplementary Table 1: List of primers used.**

| **Overlap Extension PCR Primers** | |
|---|---|
| Primer Name | Primer Sequence 5'-3' |
| hPNPLA3 I148M $^{370}$QAAA$^{373}$ - Forward | AATCTGCCATTGCGATTGTCCAGGCAGCGGCGACATGGCTTCCAGATATGCCC |
| hPNPLA3 I148M $^{370}$QAAA$^{373}$ - Reverse | GGGCATATCTGGAAGCCATGTCGCCGCTGCCTGGACAATCGCAATGGCAGATT |
| hPNPLA3 I148M $^{370}$AAEE$^{373}$ - Forward | AATCTGCCATTGCGATTGTCGCAGCGGAGGAGACATGGCTTCCAGATATGCCC |
| hPNPLA3 I148M $^{370}$AAEE$^{373}$ - Reverse | GGGCATATCTGGAAGCCATGTCTCCTCCGCTGCGACAATCGCAATGGCAGATT |
| **Cloning PNPLAs with GLucN** | |
| Primer Name | Primer Sequence 5'-3' |
| PNPLA6 - Forward | CGCGGCTAGCGCCACCATGGGGACATCGAGTCACGGGCT |
| PNPLA6 - Reverse | CGCGGGATCCGGGGCATCTGTGGCTGAGCCGGGC |
| PNPLA7 - Forward | CGCGGCTAGCGCCACCATGGAGGAAGAGAAAGATGACAGCC |
| PNPLA7 - Reverse | CGCGAAGCTTGGCCCGTCCTGGTCAGAGGAGCC |
| PNPLA9 - Forward | CGCGAAGCTTGCCACATGCAGTTCTTTGGCCGCCTGG |
| PNPLA9 - Reverse | CGCGACCGGTGGGGGTGAGAGCAGCAGCTGGATGAG |
| **hPNPLA3 C-terminus Amplification for PNPLA4 Fusion** | |
| Primer Name | Primer Sequence 5'-3' |
| PNPLA3 320 - Forward | CCAACCGGTCCCCAGGCTCGCTACAGCACT |
| PNPLA3 481 - Reverse | CCAACCGGTGGCAGACTCTTCTCTAGTGAAAAACTGGGAAA |
